# Shedding light on Gaussia Luciferase conformational space using multi-conformation enhanced sampling simulations

**DOI:** 10.64898/2026.09.18.752584

**Authors:** Berta Bori-Bru, Oriol Bárcenas, Julian Olivier Streit, Kresten Lindorff-Larsen, Ramon Crehuet

## Abstract

Gaussia luciferase (GLuc) has broad biotechnological applications owing to its bioluminescence, yet its catalytic mechanism remains poorly understood and rational engineering efforts are hindered by the absence of a structural model. This gap stems from GLuc’s highly dynamic nature, which allows it to populate a wide range of conformations rather than adopt a single, well-defined fold. Aiming to bridge this gap, in this work we employ structure predictors and conformational sampling techniques to prod GLuc’s structure. Our results indicate that structure predictors such as AlphaFold, as well as ensemble predictors such as BioEmu, fail to adequately sample this conformational space, and even extended molecular dynamics simulations do not yield converged conformational ensembles. However, enhanced sampling via PT-WTE provides a more satisfactory description of GLuc’s conformational landscape, allowing us to cluster distinct conformational basins, identify loosely defined substrate-binding pockets, and characterize the conformational changes induced by substrate binding. These insights advance our understanding of GLuc’s catalytic mechanism and inform future efforts to engineer improved variants.

## Introduction

Gaussia luciferase (GLuc) is a bioluminescent enzyme isolated from the marine copepod *Gaussia princeps.* It catalyzes the oxidative decarboxylation of coelenterazine, emitting blue light at 480 nm. GLuc is distinguished by its small size (168 amino acids, 18.2 kDa), exceptionally high bioluminescent intensity, [1]low cellular toxicity, stability across a broad pH range, and resistance to thermal shock and oxidative stress [2]. These features have established GLuc as a versatile and prominent reporter in the life sciences. However, GLuc also presents limitations, including a short half-life, restricted utility for intracellular monitoring, and high susceptibility to tissue absorption in vivo [3–5], motivating ongoing efforts to engineer GLuc variants with improved reporter properties. These efforts, however, are hampered by the absence of a structural model: unlike other well-known luciferases —such as those from Renilla, Oplophorus, or Firefly— which possess well-defined tertiary structures, GLuc lacks one.

Despite its widespread application, the structural properties of GLuc remain elusive, fueling considerable debate within the scientific community. The first reported structure, determined by NMR (PDB: 7D2O) [5], revealed a compact four-α-helix bundle connected by a disordered region. The significance of this structure was highlighted by its selection as a target in the CASP14 protein structure prediction competition [6], in which AlphaFold2 (AF2) produced the most accurate model. However, its Global Distance Test score was only 61/100—substantially lower than the average of 92/100 for other targets—underscoring GLuc’s unusual structural complexity. Addition of NMR restraints to AlphaFold predictions can reveal hidden conformations but results are sensitive to the choices made in the NMR experiment preparation and measurements [7]. Notably, Wu et al. introduced two residue mutations to enhance expression, which reduced the enzyme’s activity to less than half that of wild-type GLuc, potentially affecting its structural integrity. In response, Dijkema et al. reported an alternative structure using the wild-type sequence and a different expression system (PDB: 9FLA) [4]; this structure displayed substantially greater disorder and did not resemble 7D2O. These findings suggest that GLuc occupies a unique position between intrinsically disordered proteins (IDPs) and conventional globular proteins—exhibiting more structure than a typical IDP yet greater flexibility than a typical globular protein. As enzymes tend to be globular, GLuc joins this unfamiliar but growing family of highly disordered or fuzzy enzymes. [8,9].

Controversy also surrounds the disulfide connectivity of GLuc’s cysteine residues, with several studies suggesting conflicting bonding patterns. Because disulfide connectivity directly constrains the conformational ensemble accessible to the protein, selecting the correct configuration is a critical, unresolved parameter for structural modeling [4,5,10,11]. Further discrepancies remain regarding the location and composition of GLuc’s coelenterazine-binding site. While most publications agree on the key role of arginines R76 and R147 in enzymatic activity, the identities of other residues that delimit the binding pocket remain unclear [4,5,12]. Notably, both published NMR structures position these critical arginines far apart, suggesting that neither structure represents the active conformation of the holoenzyme [13].

The advent of deep learning predictors has revolutionized protein structure determination. Since the CASP14 competition [6], AF2 and its successor AlphaFold3 (AF3) [14] have continually improved in accuracy, contributing to a Nobel Prize [15] and stimulating a proliferation of competing methods in the protein structure prediction field [16–20]. These tools can predict the structure of proteins or protein complexes directly from their amino acid sequence or similar representations. More recently, softwares such as BioEmu [21] have taken this further by predicting ensembles of structures that emulate molecular dynamics (MD) simulations.

MD simulations provide a physics-based framework for investigating proteins with heterogeneous structural behavior. Unlike static crystallographic, NMR, or deep learning–based prediction models, MD offers a dynamic view of protein motion, capturing fluctuations and rearrangements under physiological conditions [22]. This is especially valuable for proteins with disordered or metastable regions, where a single structure may fail to capture functionally important behaviors, or when the protein interacts with a molecular partner, such as a substrate, that might affect its dynamics. Over recent decades, enhanced sampling techniques [23,24] have extended the timescales and conformational space accessible to MD, enabling characterization of rare events such as domain rearrangements, cryptic pocket formation, and folding transitions that would otherwise require prohibitively long simulations. As a result, MD has become a central tool for studying intrinsically disordered proteins (IDPs) and disordered regions, providing ensemble-based descriptions of these highly heterogeneous systems [25].

In this article, we contribute to the ongoing discussion of GLuc structure from a computational biology perspective, employing molecular dynamics and parallel-tempering well-tempered ensemble (PT-WTE), an enhanced sampling method. We first compare predictions from state-of-the-art structure predictors with available NMR structures. Then, we conduct microsecond-scale simulations of AF3 and 9FLA structures (see Methods) and compare them with BioEmu predictions. Finally, this comparison is further enhanced by employing PT-WTE simulations using both AF3 and 9FLA structures, with and without GLuc substrate, coelenterazine (CTZ), and analyzing the resulting trajectories using structure-based principal component analysis (PCA) and clustering methods to elucidate binding dynamics.

## Methods

### Structure predictors

All luciferase predictions for AF3 [14], Boltz-2x [17], and RF3 [18] were made with 10 samples, generated with 5 recycles. RF3 predictions used MSAs generated by Boltz-2x. For BioEmu [21] predictions, we sampled 500 predictions and grafted side chains with the side-chain reconstruction and MD relaxation protocols incorporated in the software.

### MD simulations

As starting structures for MD simulations, we chose the highest-scoring AF3 structure (hereafter, AF structure) and the first reported NMR structure with the associated PDB ID 9FLA (hereafter, NMR structure). We chose to use 9FLA as it has the wild-type sequence, and recent studies support its disulfide bond arrangement [11].

For a correct comparison between the simulations, we changed the disulfide bond arrangement in the AF structure using GROMACS pdb2gmx and by editing the topology file. In addition, the first 20 amino acids of the NMR structure were reconstructed using MODELLER [26], as they were not structurally determined in the original PDB.

AF and NMR simulations were performed using GROMACS 2023.3 and AMBER99SBdisp. To accelerate the simulations, we employed virtual sites on the methyl groups, which allowed us to increase the integration timestep to 4 fs. We performed minimization and equilibration in the NVT ensemble at 310K using the V-rescale thermostat. Then, we performed simulations in the NPT ensemble at 310K using the V-rescale thermostat and C-rescale barostat. We obtained 10 μs for AF simulations and 20 μs for NMR. We discarded the first 20 ns and 40 ns of AF and NMR simulations, respectively, as equilibration and extracted 500 frames for analysis. The choice of temperature was based to match the NMR measurements in [4].

### PT-WTE simulations

To enable a more comprehensive exploration of the protein’s conformational space and to analyze its interaction with coelenterazine, we employed PT-WTE simulations [27]. The PT-WTE simulation used 32 replicas over the temperature range 300-500 K. This methodology requires multiple replicas to explore the temperature space. To promote replica exchange, energy fluctuations are enhanced with a bias potential. The replica at 310 K (matching the temperature used for the rest of the simulations) was not biased, so its statistical properties could be obtained without reweighting. We employed the same GROMACS version and force field as the regular molecular dynamics. GROMACS was patched with the open-source community-developed PLUMED library [28] version 2.9.1 [29].

Given the similarities and differences between the NMR and AF structures, we decided to use a mix of both conformations to run the PT-WTE simulations. Thus, we used frames extracted from the equilibrated AF and NMR MD simulations as the initial conformations between replicas. This was done so both AF and NMR replicas were perfectly mixed, i.e., replica 1,3,5,… started from an AF conformation, and replica 2,4,6… started from an NMR conformation.

Two systems, with 1 and 2 CTZ molecules, were also simulated using the same box as the apo enzyme. We introduced the CTZ molecules at random positions within the solvent region of the simulation box. To enforce consistent exchanges between replicas involving different systems and avoid convergence issues, both AF and NMR systems had the same atoms, both in number and type, differing only in their positions. CTZ was parametrized using GAFF2 [30,31] and RESP charges, obtained from Gaussian16 [32] at B3LYP/6-311G** level of theory. The corresponding electrostatic parameters were converted to GROMACS format using ACPYPE [33].

The three PT-WTE simulations run for 500ns each. In accordance with the MD and BioEmu trajectories, we extracted 500 frames from each PT-WTE simulation, corresponding to 1 frame per nanosecond. In this case, we included the entire trajectory in the analysis for two main reasons. First, both the AF and NMR starting structures came from the end of their respective simulations. Second, PT-WTE simulations require a prior equilibration and bias phase for each replica, which is not included in the analysis trajectory.

### Analysis

To analyse the trajectories we calculated intrinsic properties of the system and performed principal component analysis (PCA) and clustering. Physical parameters like root-mean-square deviation (RMSD), root-mean-square fluctuation (RMSF), radius of gyration (Rg) and secondary structure assignment were calculated using the available functions in MDTraj [34]. For PT-WTE simulations, CTZ-Protein contacts were defined when 2 or more substrate-protein heavy atom pairs were at <5Å of distance. For the PT-WTE simulation with 2 CTZs, substrate aggregation was defined as 2 or more heavy-atom pairs from different CTZ molecules being within <5Å.

The conformations from all trajectories were analyzed using Principal Component Analysis (PCA), Dimensional Reduction Ensemble Similarity (DRES) [35,36] and an in-house-developed clustering algorithm, which will be described in detail elsewhere. PCA was performed using the Cα-Cα distance tensor from all simulations as input for scikit-learn PCA class [37]. DRES was used to assess both conformation similarity between trajectories, using PCA with 3 dimensions, and the convergence of the ensemble sampling, using window sizes of 10 frames. DRES is available in the ENCORE package from MDAnalysis [38,39]. Briefly, the clustering algorithm generates a pairwise similarity matrix for all frames across the trajectories. The structural similarity matrix is then transformed into a distance tree, which serves as the basis for clustering. Frames that are above a given similarity score (or below a given distance) are grouped into the same cluster.

## Results and discussion

### High predicted structural agreement despite low confidence

Comparison of the different predicted structures in Figure 1 shows high consistency both within each predictor’s set of models and among the different predictors, despite confidence metrics indicative of a moderately disordered protein lacking a well-defined globular fold (Figure S3A). This inter-predictor agreement is likely attributable to the fact that the training database for all these methods contains only a single deposited structure for this sequence, 7D2O: predictors converge on this specific conformation, yet their own confidence metrics correctly hint at the protein’s partially disordered nature. The effect of being in the training set probably also conditions the disulfide bond connectivity of the cysteine residues (Figure S1) and the BioEmu predictions (Figure S2), as the contact maps from the predictions are nearly identical to the one derived from the 7D2O structure.

**Figure 1:**
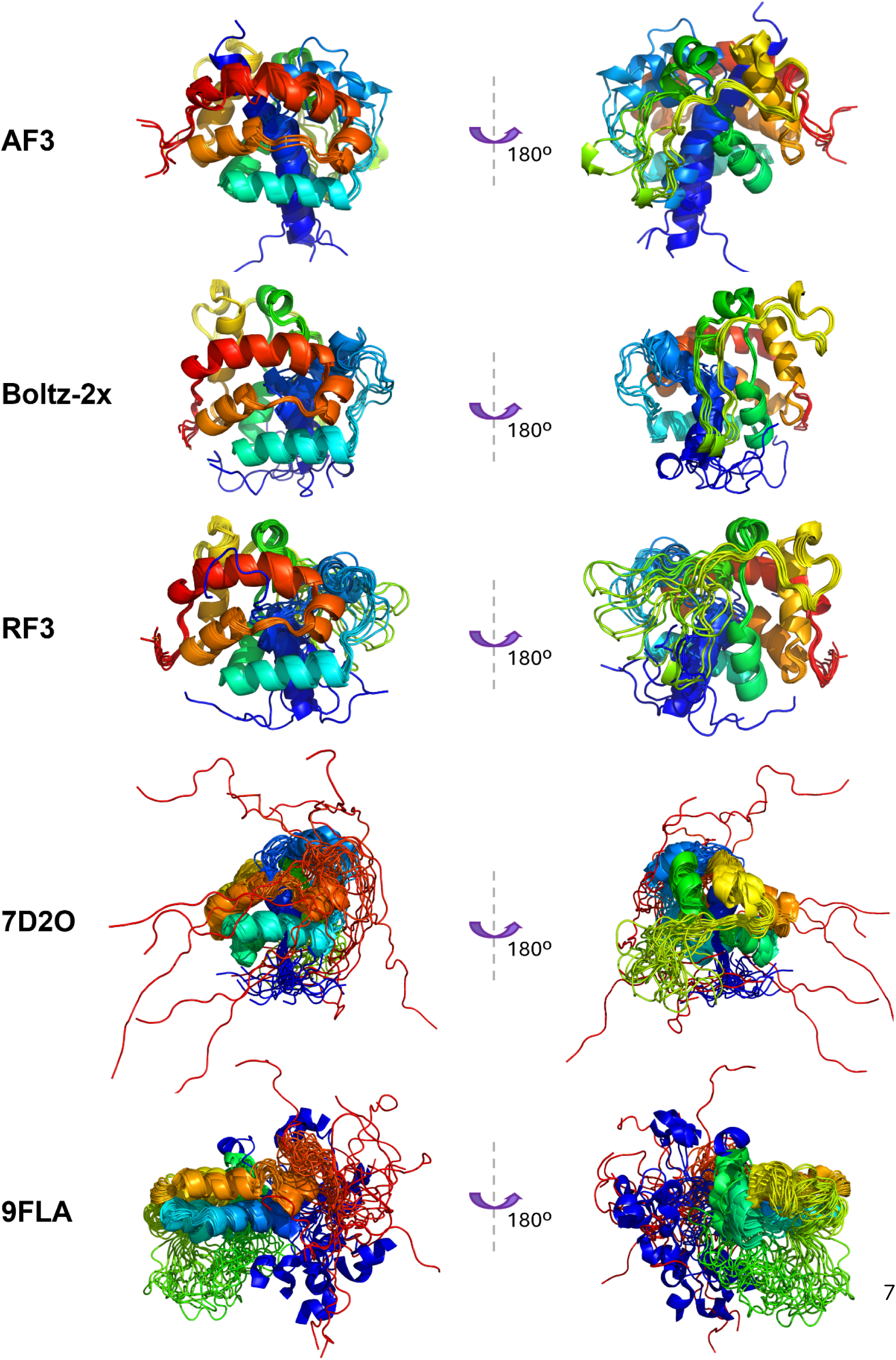
Comparison of the predicted GLuc structures with different predictors and NMR published structures, identified by their PDB id. Columns show different perspectives of the protein. Color ranges from blue for the N-terminus to red for the C-terminus.

Predictions indicate moderate-to-high agreement with 7D2O especially in the placement of the N-tail, which passes inside the protein. However, predictions show C- and N-tails with a very defined α-helix structure, which is not observed neither in 7D2O nor 9FLA. Both NMR conformations show a clear disordered behaviour in these regions. This discrepancy is apparent in Figure S3B and Figure S4, where all predictors show low pLDDT and high PAE values in the terminal and middle sections, hence highlighting that those regions might be disordered and their placement is improbable considering other protein residues.The structural ensemble and the confidence metrics position GLuc as a challenging protein to study, as it lies squarely in the middle of the order-disorder spectrum, highlighting the importance of obtaining relevant ensemble descriptors to understand structure-function relationships.

### Molecular Dynamics and BioEmu show high protein flexibility but remain trapped in their conformational basins

Given the stark differences between 9FLA NMR and the predicted structures, and the fact that GLuc confidence metrics points to a protein populating many metastable states, we decided that MD simulations would be able to describe GLuc more accurately. The intention is twofold: first, to compare how BioEmu fared against classical MD in sampling conformations for a challenging protein. Second, we wanted to know whether the starting structures of the simulations determined the conformations explored during MD, i.e., whether both NMR and AF simulations would converge to a similar ensemble. This would provide further insight into how well MD captured the protein’s underlying dynamics.

In Figure S5, we observe that classical MDs (AF and NMR) rapidly move away from the starting conformations but tend to cluster around a region in conformational space, indicating that the initial conformation is not energetically favorable and that the MDs settle into a distinct energetic basin. This is especially relevant for the NMR structure, which shifts by more than 1 nm between the first and second frames and remains in a clear energetic basin (Figure S5). Given this rapid conformational exchange and the fact that all NMR models are within an RMSD of 2Å from each other, we believe that starting from another NMR structure would not produce a much different ensemble. Moreover, AF and NMR populate different structural ensembles with very little overlap. These basins are best seen in the PCA projections in Figure 2, with no overlap between them. Although the BioEmu ensemble seems to sit between the two, this is a limitation of using only 2 principal components, as the cluster analysis reveals that BioEmu conformations are distinct from all the simulation-based methods (see Figure 4 and the associated discussion). The similarity measures of the different trajectories based on the Jensen-Shannon divergence of the principal components confirm that the sampled conformational regions of the three trajectories differ (Figure S6A) even though they seem to have locally converged (Figure (S6B). Frames are correlated to a similar degree, so that convergence seems to happen at a similar speed even though the length of the simulations is different.

**Figure 2:**
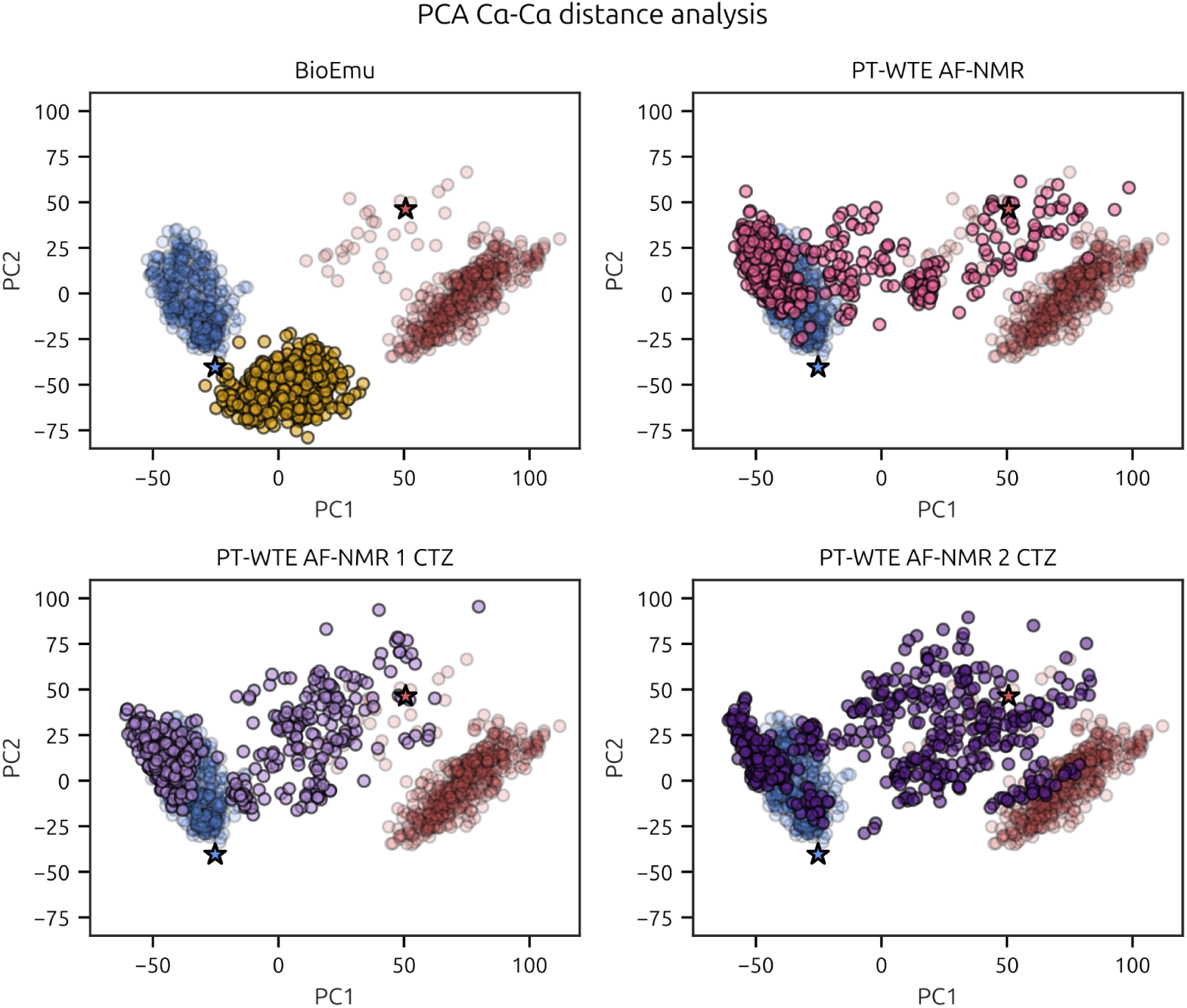
PCA analysis of all trajectories. AF (blue) and NMR (red) structures’ PCAs were anchored and compared to the other simulations. Stars indicate the starting conformations of AF and NMR. Different colors indicate different simulations. This color encoding will be used through the remaining figures of the manuscript.

To further probe the limitations of MD and better characterize the conformational ensemble, we analyzed the trajectories using both local and global metrics. RMSF, which reports local mobility, is markedly higher for AF conformations in the protein’s central helical region. In contrast, NMR and BioEmu display similarly low RMSF values, despite populating distinct regions of the structural landscape. At the global level, the radius of gyration (Rg) distributions (Figure S7) show that AF conformations average around 1.75 nm, compared to 1.8-1.9 nm for NMR, while BioEmu adopts a more compact Rg than either. Notably, analysis of the protein’s helical core reveals an unexpected trend: despite its smaller overall Rg, AF exhibits a more extended core than NMR—consistent with the greater local mobility observed by RMSF in this region.

In contrast to the previous metrics, which depend on tertiary structure, all simulations similarly reproduce the protein’s high helical propensity in the core regions, except in residues 50–60, where AF shows a lower helical tendency. The most pronounced differences arise in the C-terminal region: whereas this tail is completely disordered in the NMR structure, both AF and BioEmu trajectories show a higher helical tendency, a similar but less pronounced trend appearing in residues 70–100. Notably, it is satisfying to see this general agreement in the position and stability of helical regions extend to BioEmu, given that its conformations are generated by an AI-based method rather than sampled from a physical simulation.

To further enhance the structural insights from the molecular dynamics, we clustered the resulting structures using the algorithm described in the methodology section. We clustered the configurations from all trajectories to assess their similarity and the mixing among the clusters. Results for the enhanced sampling method and the presence of the substrate will be discussed below.

The cluster composition parallels with the PCA analysis and provides further insight into the conformational landscape, as summarized in Figure 4. Cluster 1, the major cluster, includes structures from AF and all PT-WTE trajectories. Clusters 2 and 3 are pure, containing conformations exclusively from BioEmu and NMR, respectively, indicating that these conformations remain local to their respective trajectories and are not sampled elsewhere. Clusters 4 and 5 likewise contain only AF conformations, but populate a distinct energetic basin from cluster 1, despite describing structurally similar conformations. This distinction is most apparent in cluster 4, which shows a lower Rg and narrower Rg dispersion than cluster 1 (Figure S8), consistent with closer proximity to the starting AF structure, further supported by its exclusively AF composition.

Overall, despite agreement in local secondary structure, conformational dynamics are dependent on the starting structure. Therefore we switched to enhanced-sampling simulations to better characterize the protein’s conformational landscape.

### PT-WTE is able to exchange conformations from different basins

As described in the methodology section, we performed PT-WTE simulations using initial structures coming from NMR and AF. The first step is to assess whether this methodology is adequate for the system. PT-WTE requires overlapping energies to perform replica exchange. Thus, if the structures are energetically too far apart, they are unlikely to exchange and will segregate. Results in Figures S9-S11 indicate that all replicas, regardless of whether they start from AF or NMR structures, have overlapping energetic profiles, leading to successful exchange throughout the trajectory. Although AF and NMR structures show no structural overlap during their regular MD trajectories, PT-WTE simulations successfully explore both energetic basins and, as explained later, their conformations are sensitive to the presence of the ligands. Figure 2 shows that PT-WTE simulations sample the NMR and AF conformations, but missing a region of the conformations sampled by the NMR MD trajectory. The time evolution of these PCs (Figure S12) reveals that there is not a drift towards the NMR or the AF basins, confirming that both basins are on a par in energetic accessibility. DRES analysis (Figure S6A) indicates that PT-WTE simulations sample a similar simulation space, distinct from NMR and BioEmu and, to a lesser extent, AF, as shown by the PCAs.

The radius of gyration (Figure S7) shows a distribution intermediate between AF and NMR, however the distribution of the Rg of the core is much wider than that of NMR, even broader than AF, proving that PT-WTE is sampling more conformations than the mere mixture of AF and NMR plain MD trajectories. As will be discussed below, these belong to a different cluster than those sampled by the NMR MD trajectory. This can also be seen in the RMSF plots in Figure 3A, which show large fluctuation values for the central region spanning residues 80–100, closer to the AF distribution. Nonetheless, PT-WTE fluctuations are higher than AF also in the region 60-80 and in the tails. As with previous simulations, there is agreement in the amount of helical secondary structures. Interestingly BioEmu is the method that better reproduces the PT-WTE results in terms of helicity (Figure S13) as well as ensemble similarity measured with the Jensen-Shannon divergence (Figure S6A). However this also highlights an intrinsic limitation of BioEmu: its inability to capture environmental effects. Indeed the BioEmu ensemble is most similar to the PT-WTE containing two substrate molecules even though it should be more similar to the ensemble without substrate.

**Figure 3:**
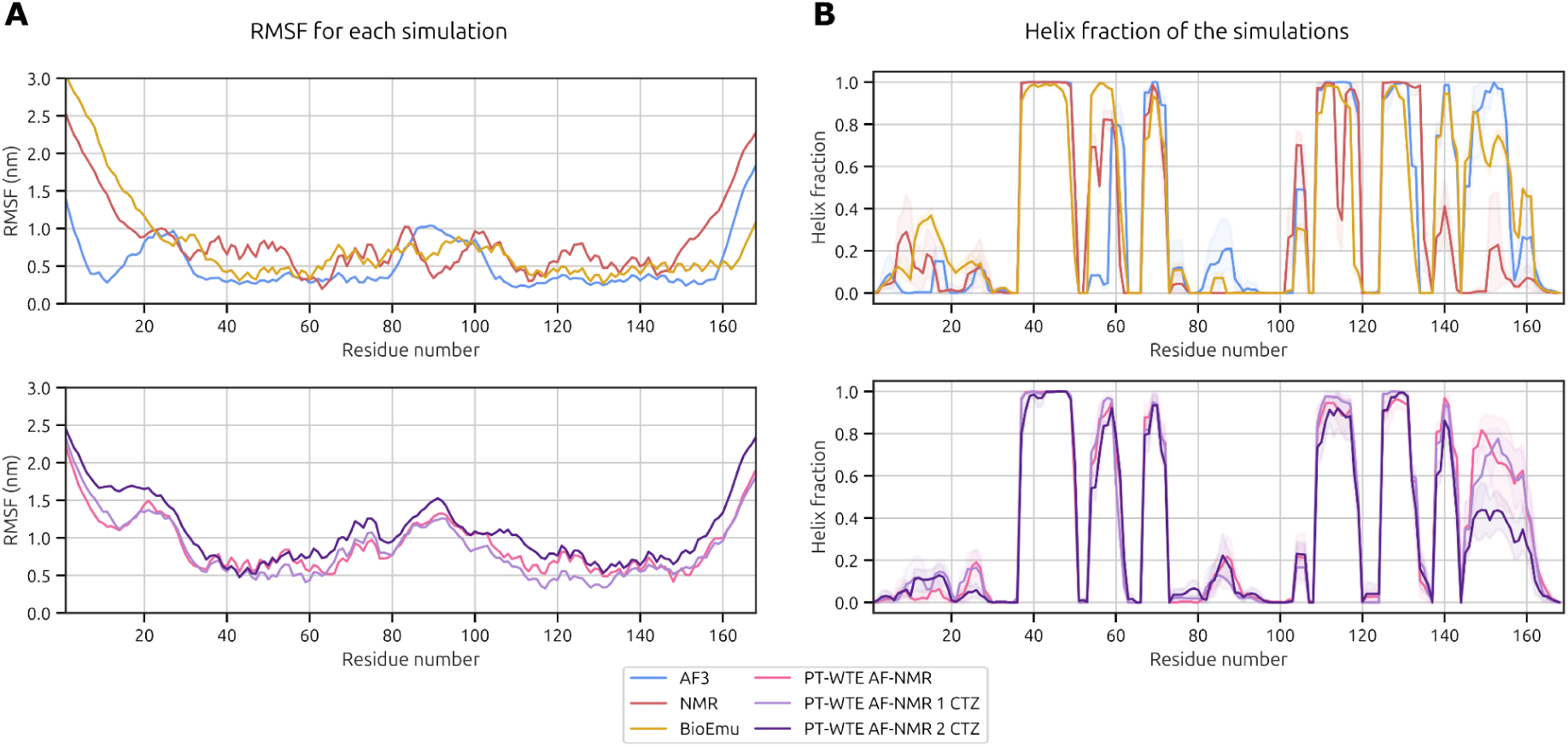
Per-residue A) RMSF and B) helical propensity for the simulations. At the top, AF, NMR, and BioEmu. At the bottom, PT-WTE simulations.

**Figure 4.**
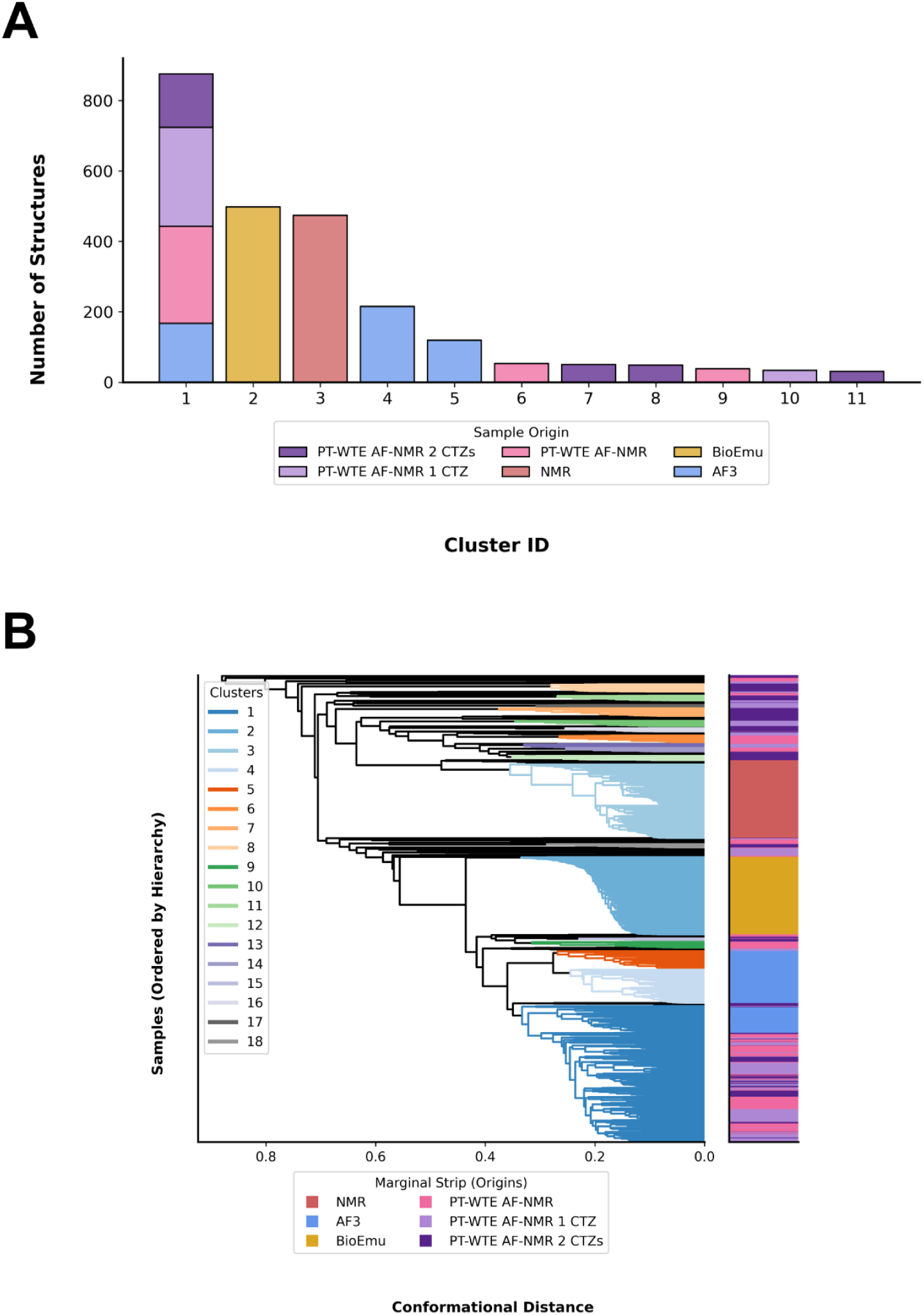
Clustering dendrogram and cluster composition analysis. A) Two color schemes are used to distinguish which structures co-cluster and which belong to the same simulation system. At the bottom, a legend lists the names of the simulated systems. These colors are also shown on the right strip: the horizontal regions covered by each color correspond to different simulation systems. On the other hand, on the left legend, we can observe colors corresponding to clusters, ordered by size (1 being the biggest cluster and 18 the smallest). These colors are shown in the dendrogram branches. One example is cluster 1 (dark blue, bottom branches), which contains structures from AF3 and PT-WTE simulations, as indicated by the colors in the right strip. B) Cluster composition analysis of structures coming from molecular dynamics simulations. Each bar corresponds to an individual cluster ID, with colors corresponding to their system of origin.

PT-WTE extended conformational sampling is confirmed by cluster analysis. The PT-WTE trajectory populates cluster 1, where AF conformations also reside. It does not contribute to the BioEmu cluster (cluster 2) and NMR cluster (cluster 3) and smaller AF clusters (clusters 4 and 5). This suggests that these conformations are metastable. In fact, many of the sampled conformations are close to the experimental 9FLA NMR structure, whereas the molecular dynamics trajectories drifted away from this structure and remained trapped in a separate basin (see also Figure 2). Indeed the 500 frames of PT-WTE trajectory are spread along 8 clusters, with more than 10 members, whereas the AF and NMR trajectories populate only 3 and 1 cluster, respectively. Thus, it seems clear that PT-WTE can explore many different conformational basins, in contrast to regular MD or BioEmu.

Taken together, these results indicate that PT-WTE achieves broader and more thorough sampling of the conformational landscape than conventional MD, capturing conformations beyond those accessible to either AF or NMR trajectories alone, while remaining consistent with both in terms of secondary structure content. Having established that PT-WTE provides a more complete representation of GLuc’s conformational ensemble, we next investigated how the coelenterazine substrate binds across this expanded set of disordered conformations.

### Addition of GLuc substrate changes the energy landscape

Given the pronounced conformational flexibility of GLuc established above, we next examined how the enzyme engages its substrate, coelenterazine (CTZ), and whether ligand binding in turn reshapes the accessible conformational space. Previous reports suggest that GLuc harbors two symmetrical binding sites [40] and exhibits substrate cooperativity [11,41,42]; to probe this possibility, we performed simulations with either one or two CTZ molecules bound. For consistency with the apo simulations, we retained the same simulation box size, which results in an effective CTZ concentration of 4.3 mM in the two-ligand system. This concentration substantially exceeds the reported aqueous solubility of CTZ (0.16 mM), arising from its aromatic, uncharged character at neutral pH, raising the possibility of ligand aggregation artifacts, which we monitored throughout the trajectories (Figure S14). The results indicate that coelenterazine was aggregated 58% of the time, but interaction frequency with GLuc was similar in both cases, 81% of contacting time for aggregated CTZs and 84% for non-aggregated CTZ, meaning that aggregation does not prevent binding.

The presence of CTZ molecules causes an increase of the Rg (Figure S7). This increase is gradual from zero to one, and then to two CTZ molecules. Since the core Rg is not significantly affected by the presence of CTZ, this suggests that the ligand prevents the collapse of the protein’s terminal regions into the core. This can occur either through direct interaction with the terminal regions or by precluding interactions between the core and these regions. In the following section we explain that it is due to both phenomena, with predominant precluding interactions.

The effect of the CTZ on GLuc’s RMSF and secondary structure is minimal with the addition of one CTZ molecule. The addition of two CTZs, however, slightly increases overall RMSF and lowers C-terminal helicity, due to interaction with the substrate, as we will develop in further sections. This is consistent with the reported catalytic need of GLuc’s terminal regions, whose deletion significantly lowered the enzyme’s activity [4].

If we focus on the substrate simulations, except for cluster 1, no other clusters contain conformations from the apo simulations. This indicates that simulations containing CTZ explore some unique regions of the conformational space. However, most conformations remain in cluster 1, indicating that CTZ does not substantially alter the conformational landscape of GLuc. This similarity can also be seen in the lower Jensen-Shannon divergence among all PT-WTE simulations, compared to other sampling methods. The persistence of a dynamic enzyme even in the presence of the substrate, not leading to the stabilization of a single structure, is consistent with NMR results probing the conformational equilibrium in the presence of the oxidized CTZ, the reaction product of GLuc [4].

### Clustering reveals the substrate binding conformations

To further investigate the relationship between GLuc conformation and its interaction with coelenterazine, we performed structural clustering using only PT-WTE systems. The resulting six clusters (Figure 5) closely resemble the previous clustering composition. The first two clusters contain a mixture of systems, while the remaining clusters are homogeneous. Cluster 1 displays a largely homogeneous contact profile, with a shallow contact frequency distribution across all protein residues (Figure 6). PCA data indicate that cluster 1 is the most AF-like structure (Figure S15). In contrast, cluster 2 exhibits contacts primarily involving the N-terminus and, to a lesser extent, the central and C-terminal regions. Interestingly, this cluster contains conformations compatible with the PCAs of NMR structures.

**Figure 5:**
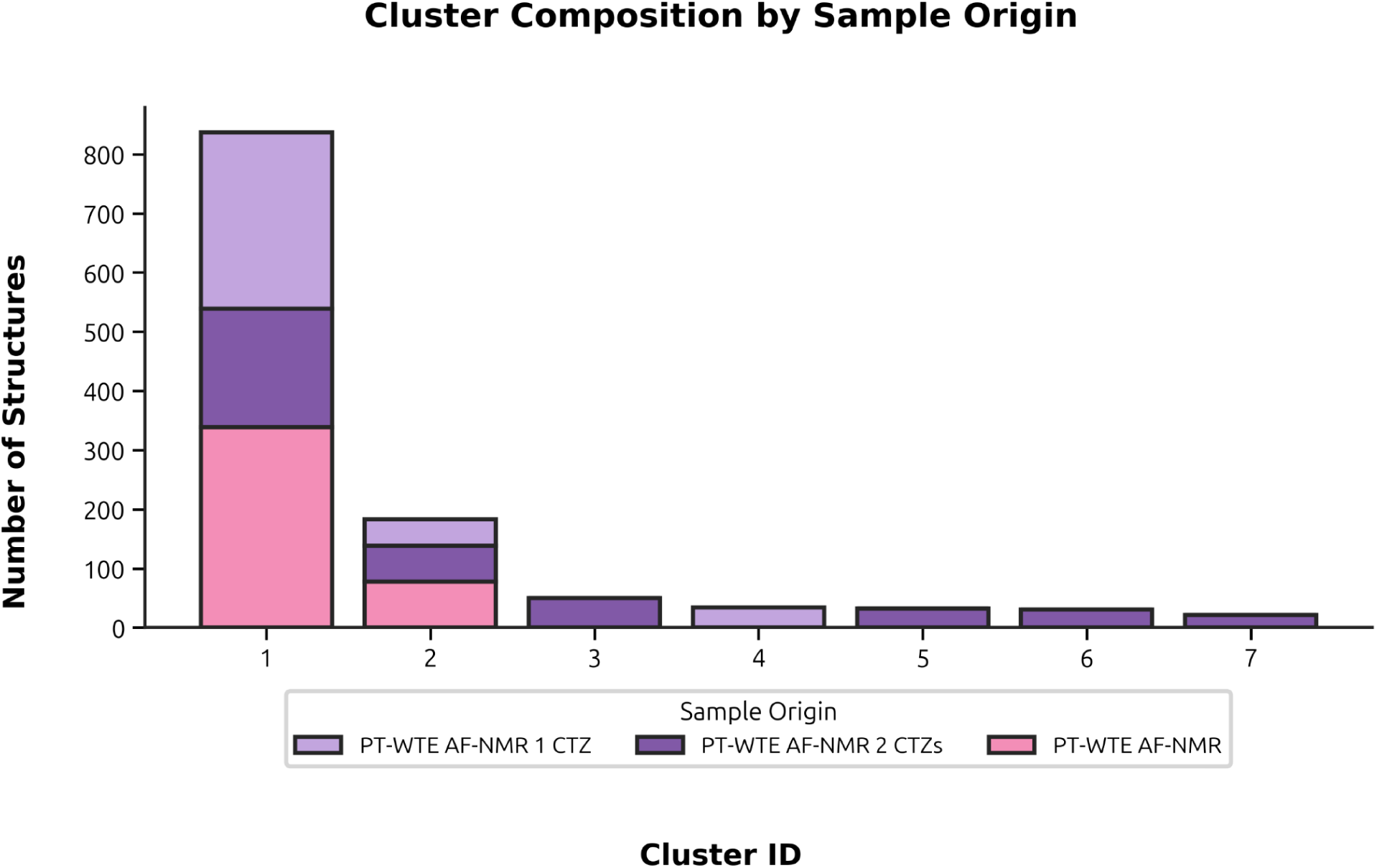
Cluster composition of PT-WTE simulations. Each bar corresponds to an individual cluster ID, with colors corresponding to their system of origin.

**Figure 6:**
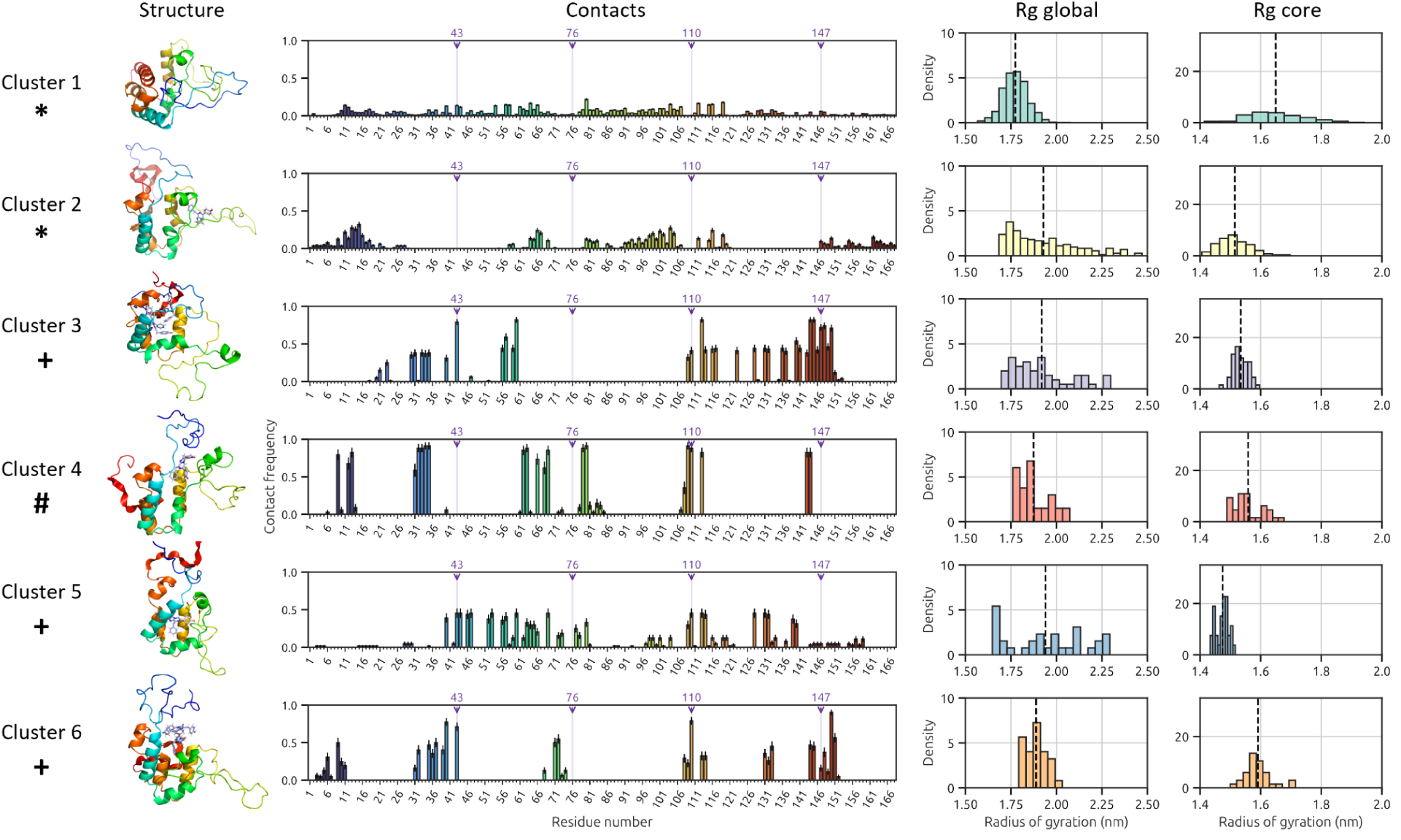
Structure and frequency of contacting residues and Rg per cluster. Shown structures are representatives of the cluster conformations. *, + and # indicate that the cluster is formed by structures from simulations with 0,1 and 2 CTZs (*), 2 CTZs (+) and 1 CTZ (#).

The absence of a defined binding region in both clusters suggests that binding may result from an induced-fit phenomenon [12], in which clusters 1 and 2 include conformations where the protein lacks a well-defined binding pocket that is subsequently induced by interaction with coelenterazine. This interpretation is further supported by the presence of structures from the substrate-free system in both clusters 1 and 2, and as we have mentioned, consistent with NMR results [4].

In contrast to clusters 1 and 2, other clusters exhibit a more defined binding pocket. The regions involved include residues 30-80 and 100-150, with some variation between clusters, corresponding to the most structured parts of the protein. In certain clusters, N- and C-terminus involvement are also observed. The results indicate that residue M43 exhibits the highest number of contacts throughout the simulations, followed by M110 (Figure S16). These findings are consistent with those reported by Borum *et al*. [11], where M43 and M110 are oxidized as a result of CTZ catalysis and thus, these residues must be near the substrate in the reaction process. Additional residues with high contact frequency include L60, F113 and T150, which may stabilize coelenterazine.

R76 and R147 are the most agreed on residues that contribute to CTZ binding in GLuc, however, our simulations do not display significantly enriched contact frequencies overall on these residues. Although clusters 3 and 6 demonstrate frequent contacts with R147, this pattern is not observed in other clusters. These observations suggest that further protein rearrangement may be required for both arginine residues to simultaneously contact the ligand.

Wu *et al.* identified additional residues that contact the ligand [5]. In the present analysis, residues A13, N17, A19, and T20, located in the N-terminus, do not exhibit high contact frequency. For other relevant residues in the central region (N46, A47, R48, F72, I73, R76, and T79), only cluster 5 displays the closest correspondence. Examination of relevant C-terminal residues (V134, R147, F151, and A152) reveals regional agreement, particularly in clusters 3,6 and to a lesser extent in 5. Overall, these findings are consistent with the binding regions identified by Wu *et al.*; however, the pronounced flexibility of GLuc complicates identifying a single, well-defined binding site or a specific set of binding residues.

Dijkema *et al.* proposed E44, R54, Y80, and D168 as relevant residues based on alanine-scanning experiments, while K70 and W143 were identified from NOE data [4]. In our simulations, we only observe binding interactions for residues E44 in cluster 5, Y80 in clusters 4 and 5, and W143 in clusters 3 and 4. Other residues do not exhibit pronounced binding interactions.

In previous sections we hypothesised that the ligand prevented the collapse of terminal regions into the protein core by either interacting directly with the protein tails or by occluding the core. Hence, we analysed the changes in global and core Rg of GLuc of each cluster to assess the effect of binding in tail mobility. In Figure 6 we found that clusters 2, 3 and 5 have the widest range in global Rg, similar to that presented in Figure S7. However, their core Rgs were much narrower. Comparing these results with the contact distributions of clusters 2, 3 and 5, we observe some contributions of the C-tail. This means that the C-terminus interacts slightly with CTZ while bound to the core. This phenomenon, in its turn, hinders protein tail collapse. Interestingly, cluster 5 has the lowest mean core Rg, similar to plain NMR simulations. For clusters 4 and 6, we observe contacts with the N-terminus, which could explain their narrower global Rgs.

Considering the role of CTZ aggregation in binding (Table S1), we observe that aggregation is relevant only in cluster 6, which is the only cluster with both high aggregation and 2-CTZ contacting fractions. We are unsure if this cluster could be an artifact due to the high CTZ concentration in the simulations. In contrast, cluster 5 shows no CTZ aggregation but does show contacts with 2-CTZ molecules. This is coherent with observed substrate cooperativity [41].

Overall, the results are broadly consistent with previously reported binding regions. Several residues identified in earlier studies are also observed in these simulations, although their contact frequencies vary substantially between clusters. GLuc’s pronounced flexibility, combined with structural differences among clusters, complicates the definition of a single binding site or a unique set of ligand-contacting residues. Furthermore, experimental data may capture regions that contact coelenterazine at different times rather than simultaneously.

## Conclusions

In this paper we observed that classical MDs starting from AF and NMR systems, although containing the same protein sequence, explored different local energy basins, leading to disparate conclusions. BioEmu also sampled one conformational basin, again different from those obtained through MD, but showed high agreement with the secondary structure of GLuc, even in partially helical regions. Hence, to explore systems with a complex tertiary structure like GLuc is advisable to use enhanced sampling methods. Our approach of using PT-WTE starting from two different conformations seems to be a valid strategy for sampling protein conformations from different energetic minima. It explores many more conformational basins beyond the starting ones, where MD remains mostly trapped.

As important as it is to correctly sample the conformational space, it is also crucial to correctly analyse the different structures that form that space. We found that our in-house clustering algorithm allowed for a more fine-grained description of GLuc’s conformations than using PCA. Specially, this allowed us to identify the different binding modes in PT-WTE CTZ simulations and define their contact profiles.

The addition of CTZ in the simulations significantly impacted the resulting conformations, which clustered differently from those in the AF and NMR simulations. The physical properties of these structures, like Rg, are also unique. From these conformations, we determined that M43 and M110 form the binding pocket of GLuc, in agreement with experimental studies. However, we found low agreement with R76 and R147 as interactors with CTZ, which might mean that we need more simulation time until convergence of the results to converge or that these residues are not involved in binding but play a crucial structural role.

We also found it quite difficult to identify a single cluster that satisfied all the binding contacts described in the literature. However, cluster 5 had the most agreement in both the Wu et al.[5] and Dijkema et al.[4] propositions. In addition, it had the most distinct, NMR-like Rg distribution and showed no CTZ aggregation, while still binding 2 molecules of CTZ. This conformation seems promising for further studies.

Finally, our research opens the door to additional experiments to supplement GLuc’s structural understanding. Further PT-WTE simulations can be performed by mixing the Wu et al. and Dijkema et al. structures to gain insights into their relative energetic stabilities. They could also reveal more about the binding modes of GLuc with CTZ. Similarly, some analyses could be repeated with different disulfide bonds arrangements. As this is a fundamental parameter in MDs, we expect different results from those presented, but with promising implications for the field. Lastly, we could study the catalytic properties of different conformational basins to further investigate the reaction mechanism of GLuc. The design of sequences that stabilize the catalytic basins could lead to engineered GLuc variants with improved applicability.

## Data Availability

The Zenodo repository with DOI 10.5281/zenodo.22765671 contains files for the 6 trajectories discussed in this paper.

## Acknowledgements

The authors thank Jakob Winther for thoughtful discussions during the development of the paper. Claude model Sonnet-5 was used to suggest alternative phrasings of some sentences of the manuscript, but never adding to or changing the meaning of the original text by the authors.

