## Supplementary figures for "Shedding light on Gaussia Luciferase conformational space using multi-conformation enhanced sampling simulations"

### Supporting information

Berta Bori-Bru[‡,1,2], Oriol Bárcenas[‡,1,3], Julian Olivier Streit[4], Kresten Lindorff-Larsen[4], Ramon Crehuet[1]

[1]Institute for Advanced Chemistry of Catalonia (IQAC) - CSIC, Barcelona, Spain

[2]Doctoral Programme in Theoretical Chemistry and Computational Modelling, Universitat de Barcelona, Barcelona, Spain

[3]Institute of Biotechnology and Biomedicine (IBB) and Department of Biochemistry and Molecular Biology, Autonomous University of Barcelona, Bellaterra, Barcelona, Spain

[4]Linderstrøm-Lang Centre for Protein Science, Department of Biology, University of Copenhagen, Copenhagen N, Denmark

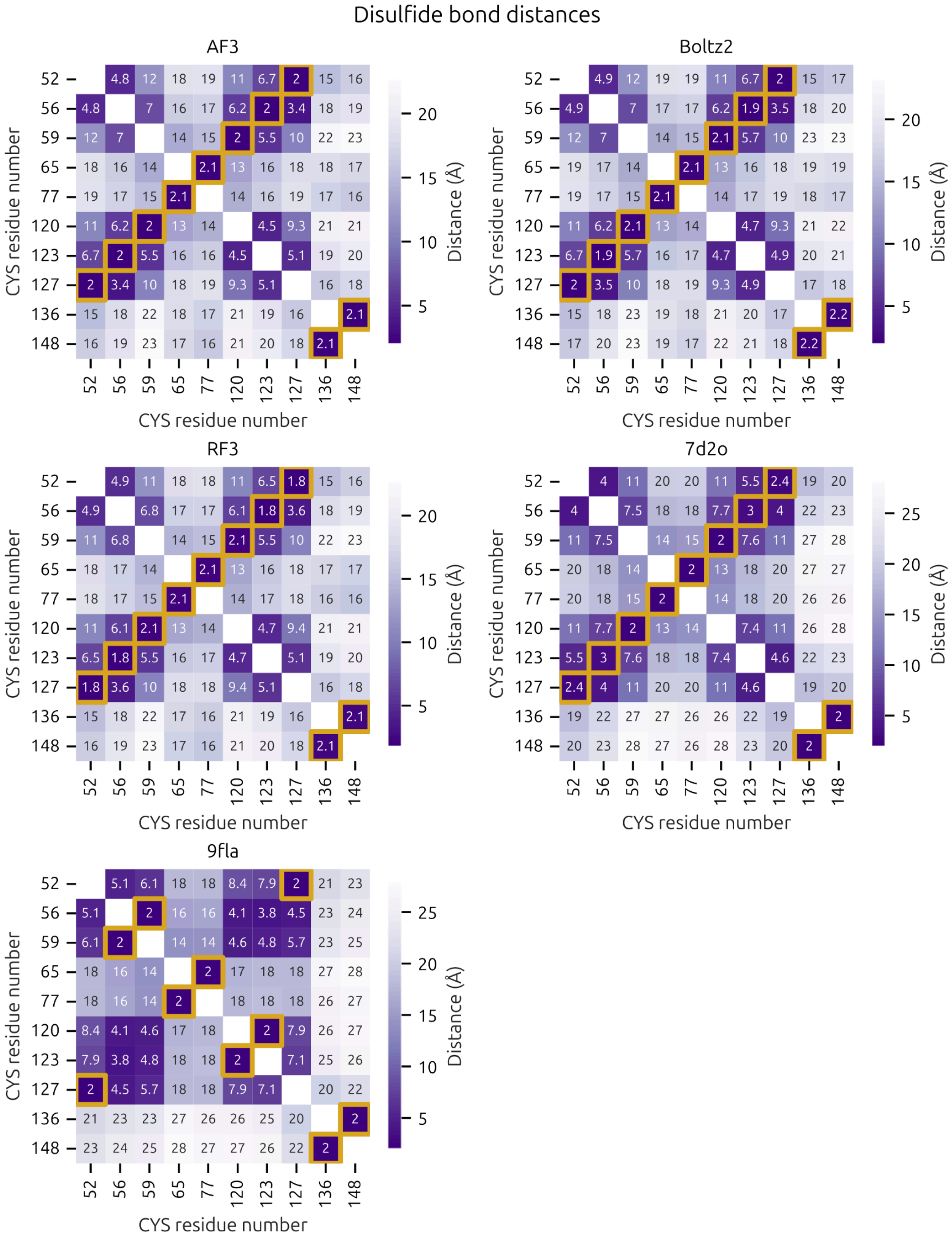


Figure S1: Disulfide bridge distance map for AF, Boltz-2x and RF3 predictions and NMR structures. Yellow squares mark the lowest distance for each residue pair, indicating the most probable disulfide bond. Distances are calculated as the mean of 10 predicted samples.

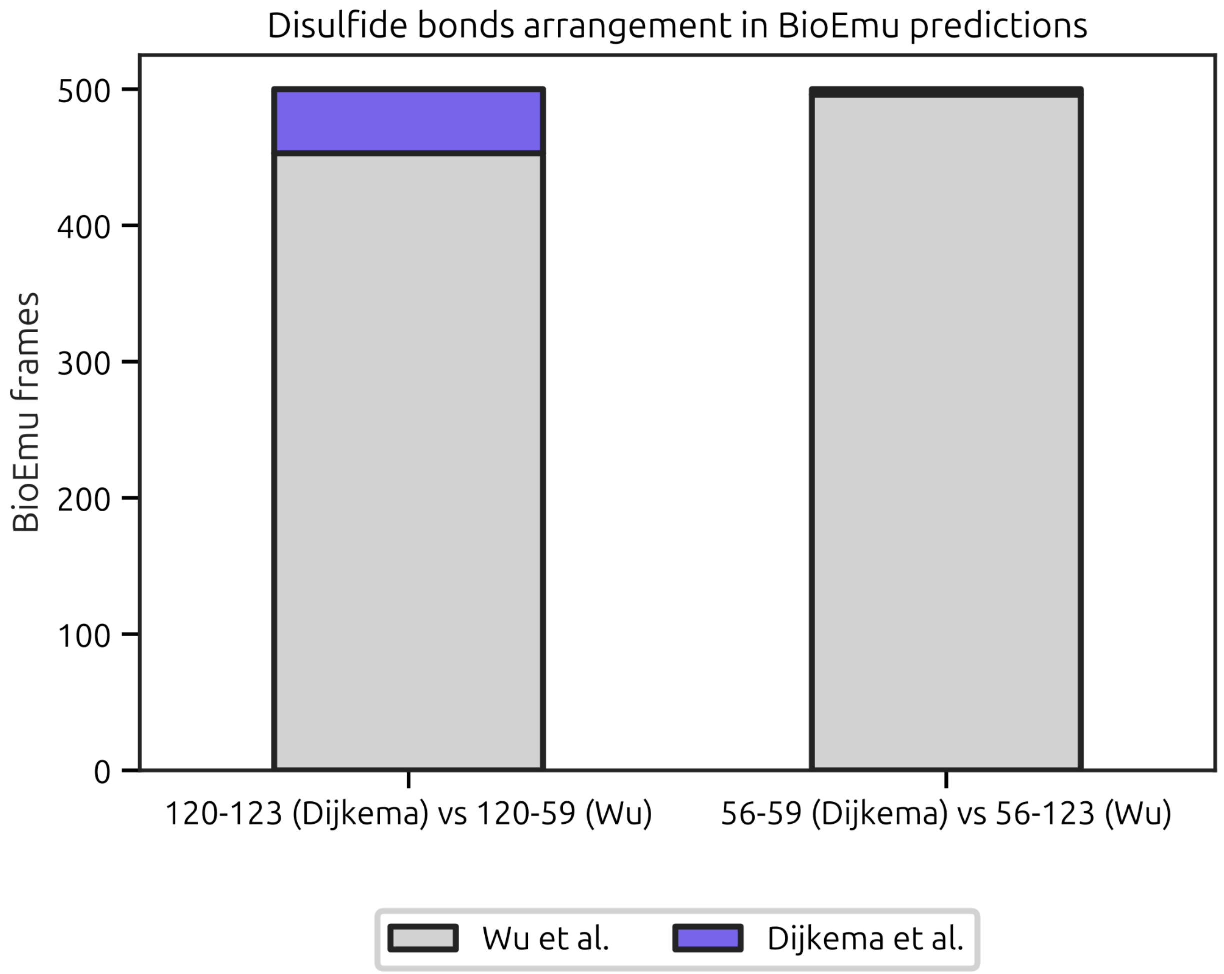


Figure S2: Disulfide bond plausibility for BioEmu predictions, identified as the number of frames that predict a structure in agreement with Wu et al. or Dijkema et al. connectivity.

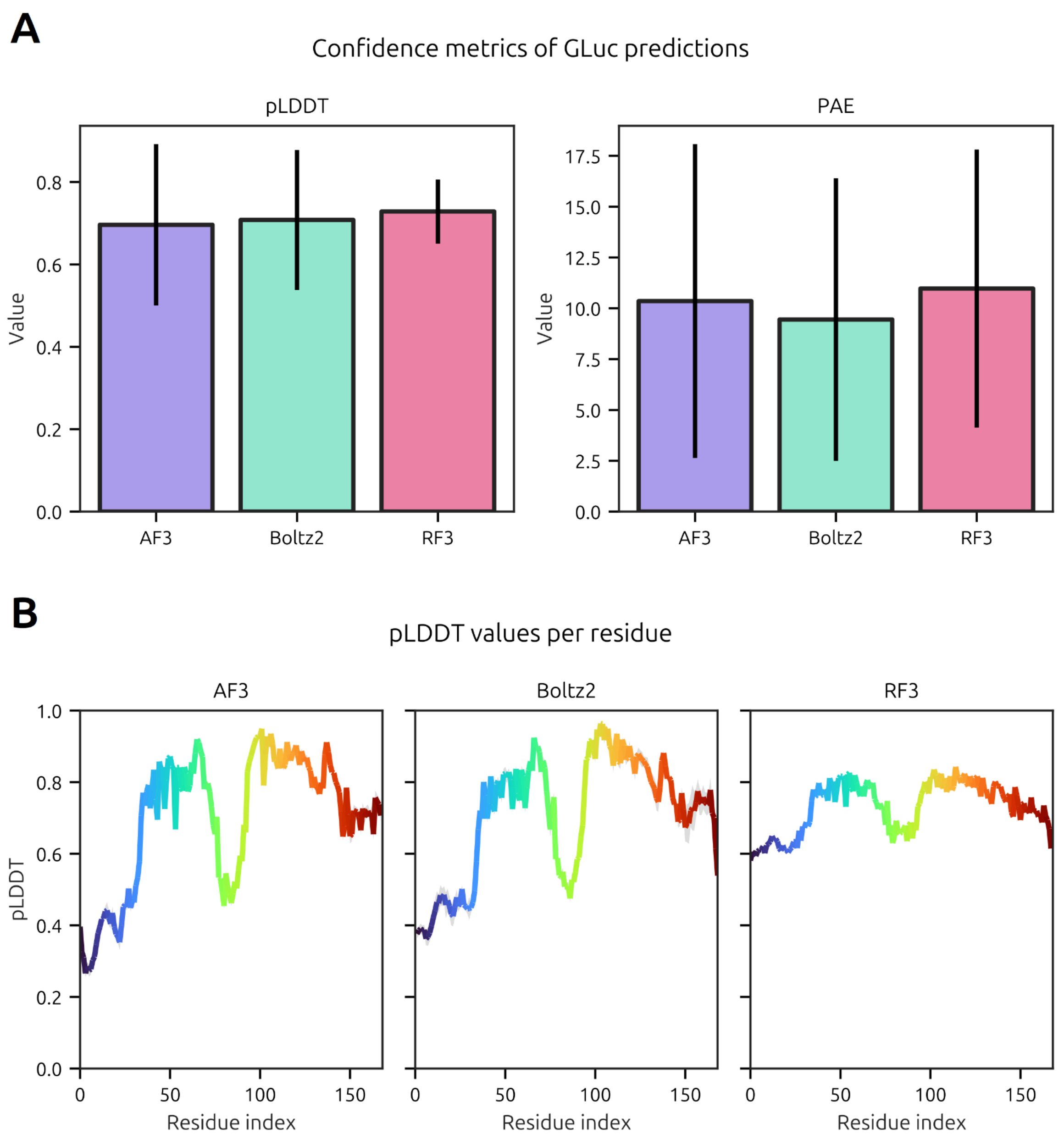


Figure S3: Confidence metrics of the predictions for GLuc performed by AF3, Boltz-2x and RF3. The error is given considering 10 different samples. (A) Average pLDDT (arbitrary units) and PAE (Å) values for the predictors. (B) Per residue pLDDT values for the predictors. Error is so small that cannot be appreciated for AF3 and RF3. Same color ranges as in Figure 1.

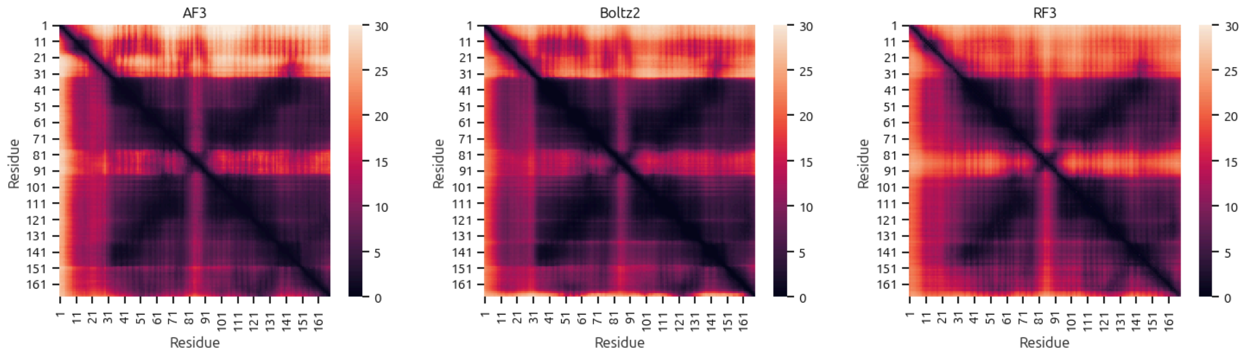


Figure S4: PAE matrixes (Å) for AF3, Boltz-2x and RF3. Obtained through the mean of the PAE matrixes of 10 predicted samples.

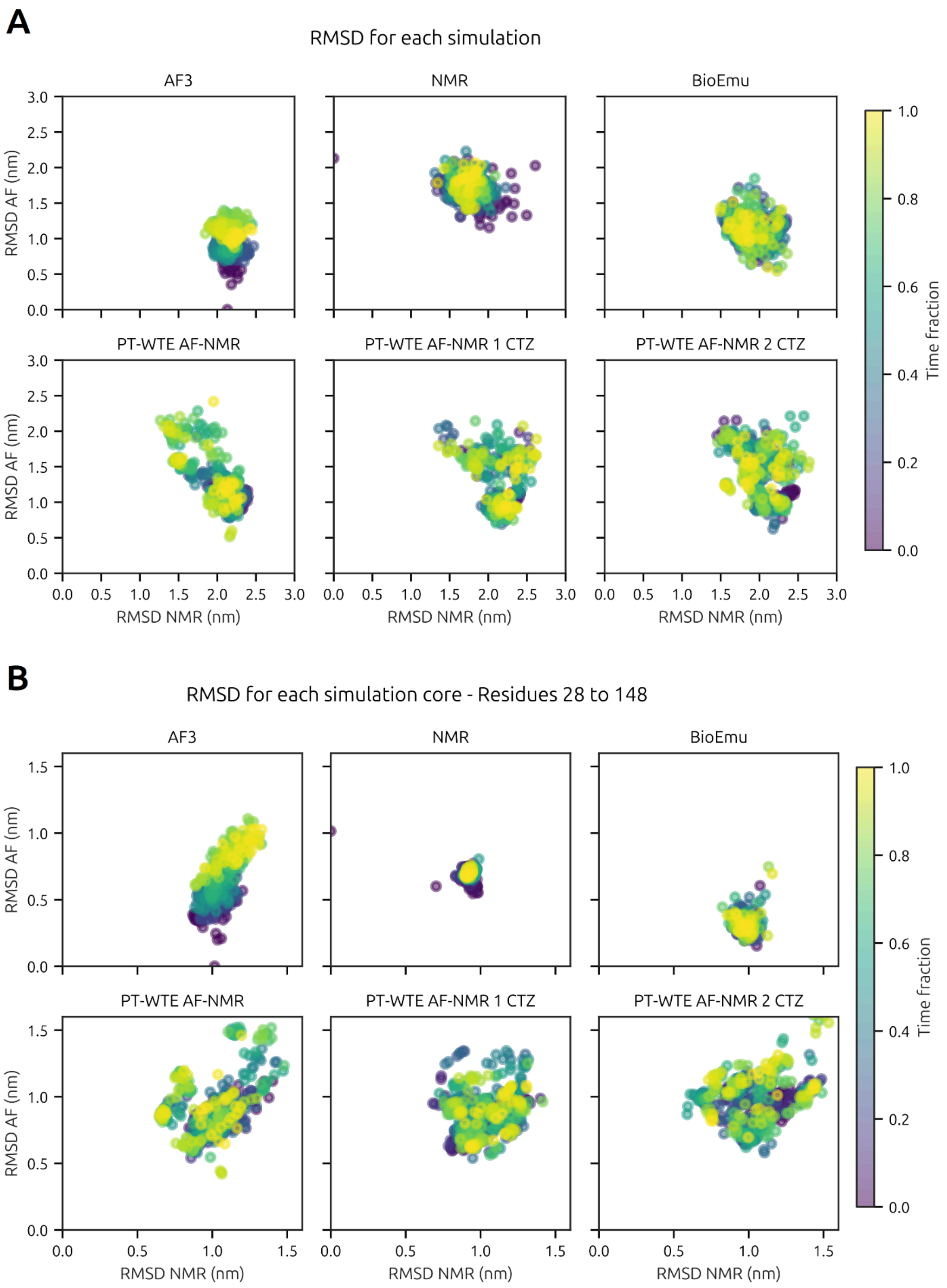


Figure S5: RMSD of the simulations A) considering all residues B) considering only residues 28-148, discarding terminal regions.

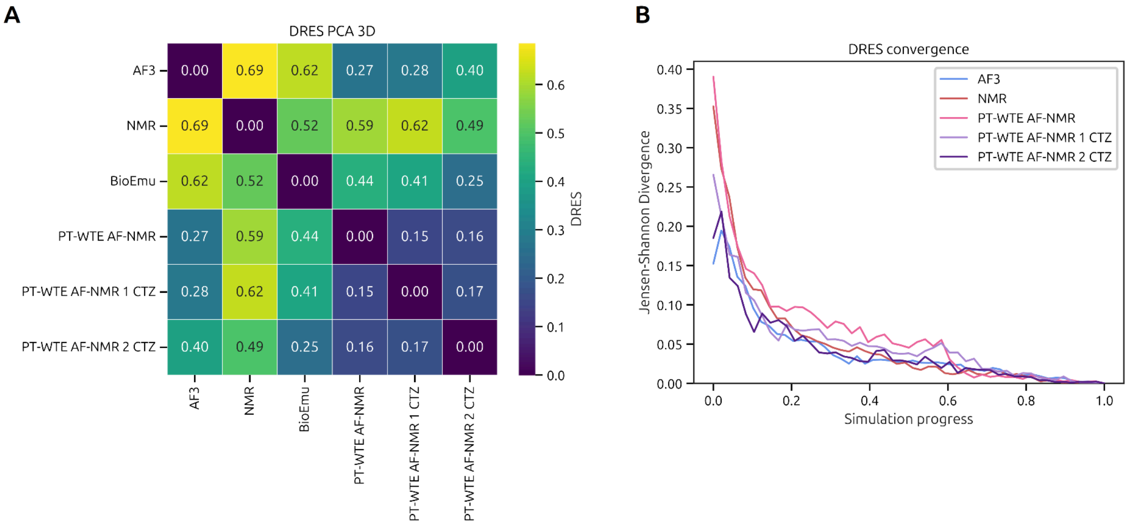


Figure S6: Dimensionality Reduction Ensemble Similarity (DRES) analysis of all simulations. A) Similarity of the ensembles sampled in each simulation, using PCAs of 3 dimensions as the dimensity reduction algorithm. The lower the value of DRES, the more similar the compared ensembles. B) Convergence of sampling in the conformational space. The faster the rate each line reaches zero, the faster the simulation samples a known conformational space. BioEmu is not shown as frames do not correspond to physical time.

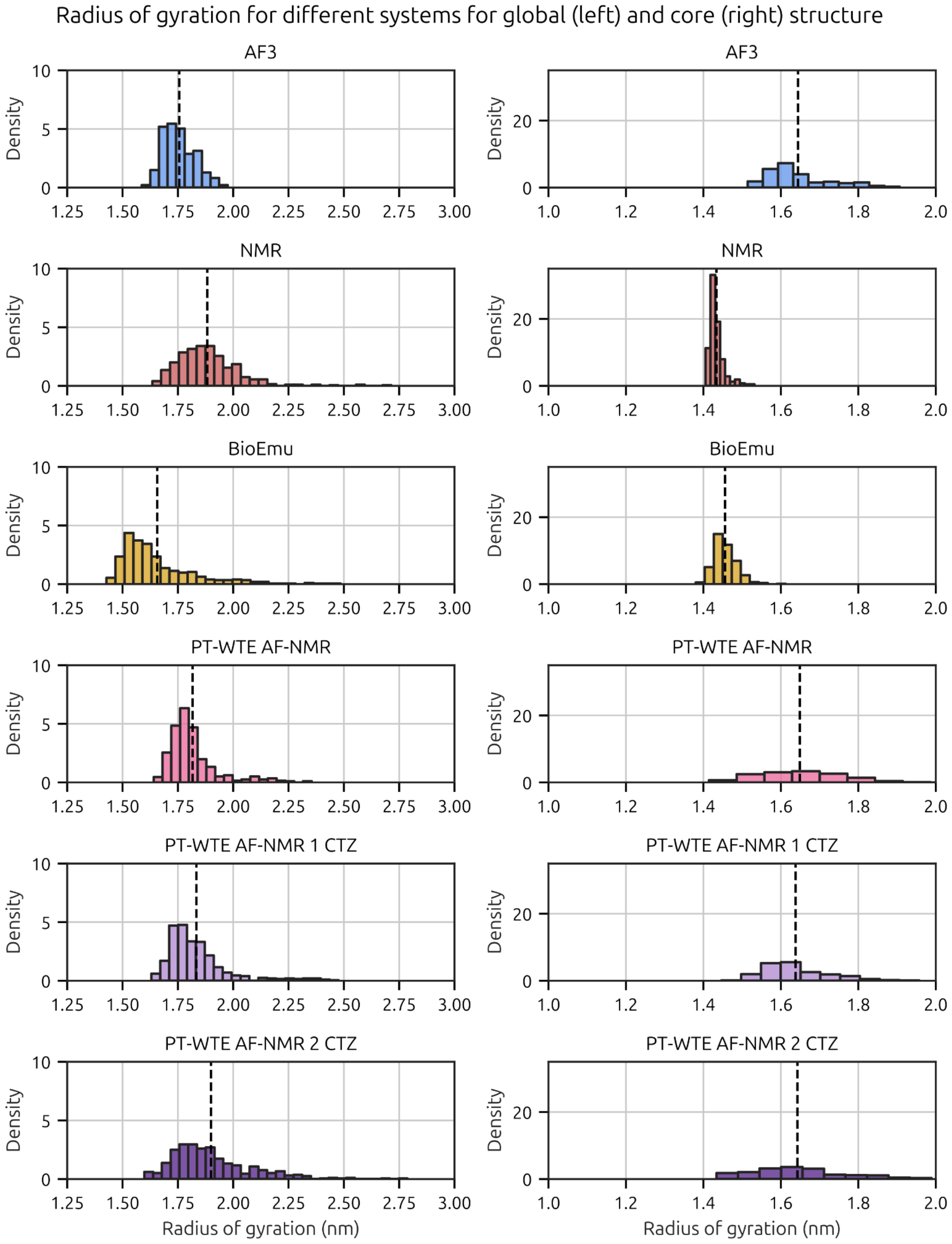


Figure S7: Radius of gyration for the different simulations. Global indicates all residues were taken into account and core indicates only residues 28-148 were considered in the analysis.

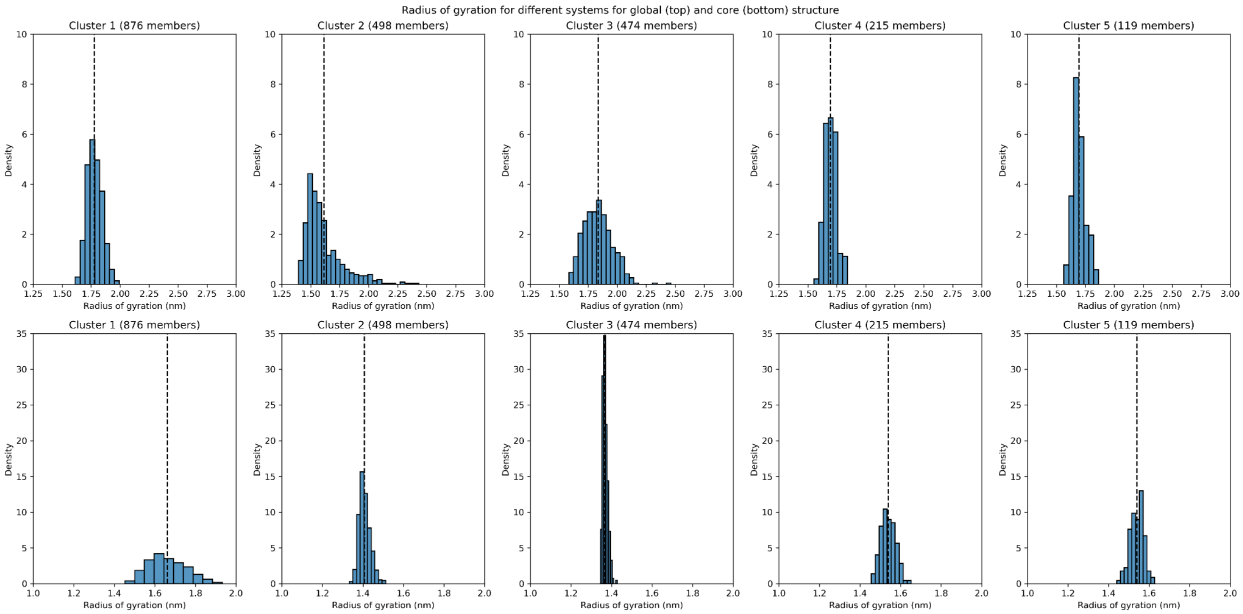


Figure S8. Rg analysis of the first 5 clusters, focusing on the overall Rg (top row) or the core's Rg (bottom row).

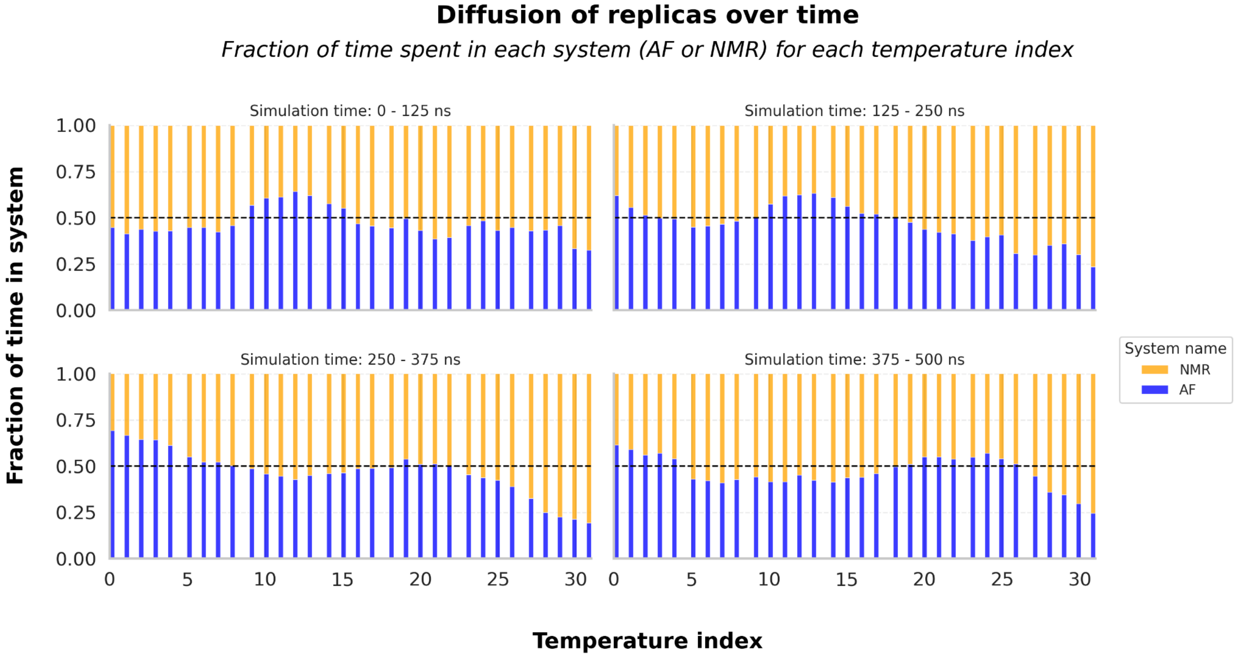


Figure S9. Analysis of the diffusion of each system for each temperature index in each simulation quarter. Replicas with NMR as the starting system are colored yellow, whereas replicas with AF as the starting system are colored blue. A horizontal dashed line marks a fraction of 0.5, indicating perfect mixing.

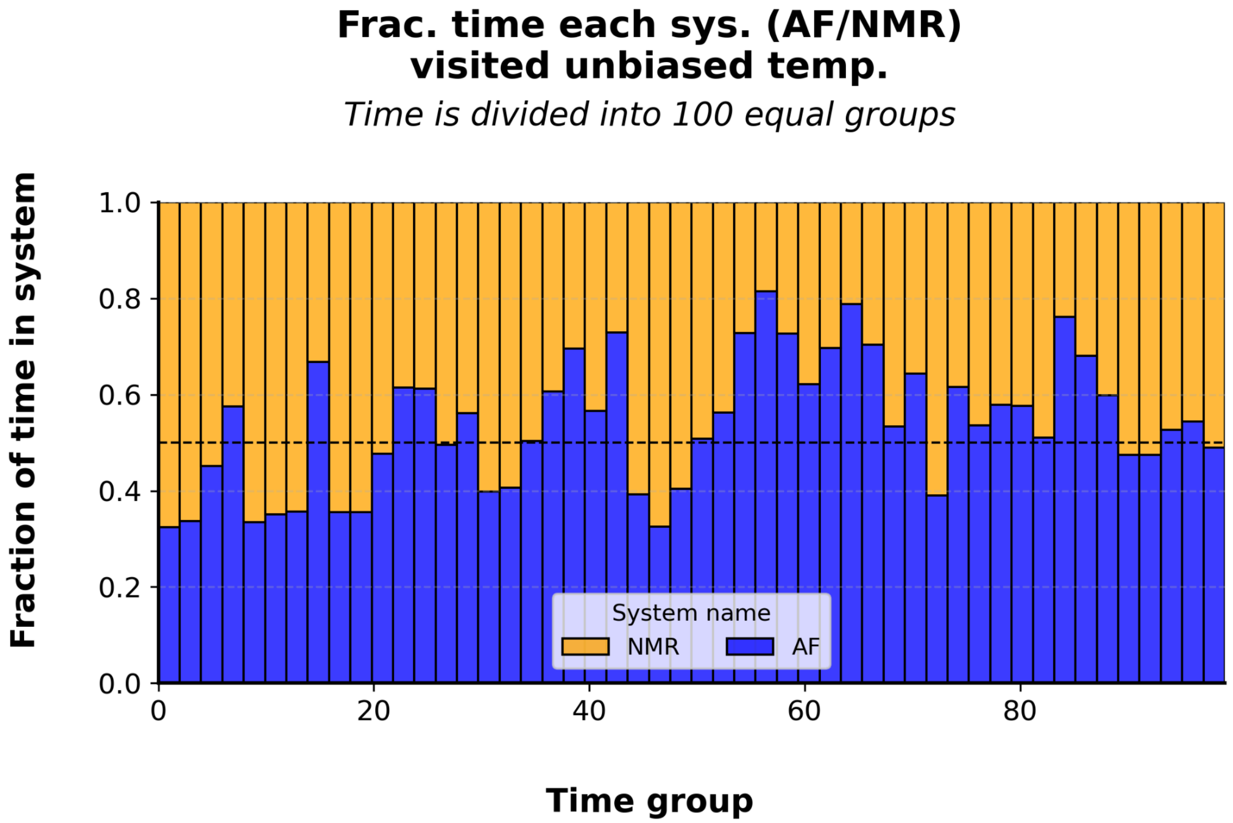


Figure S10. Analysis of the fraction of time that replicas containing NMR and AF as their starting structures are simulated at the unbiased temperature, which is used for analysis. Replicas with NMR as the starting system are colored yellow, whereas replicas with AF as the starting system are colored blue. A horizontal dashed line marks a fraction of 0.5, indicating perfect mixing.

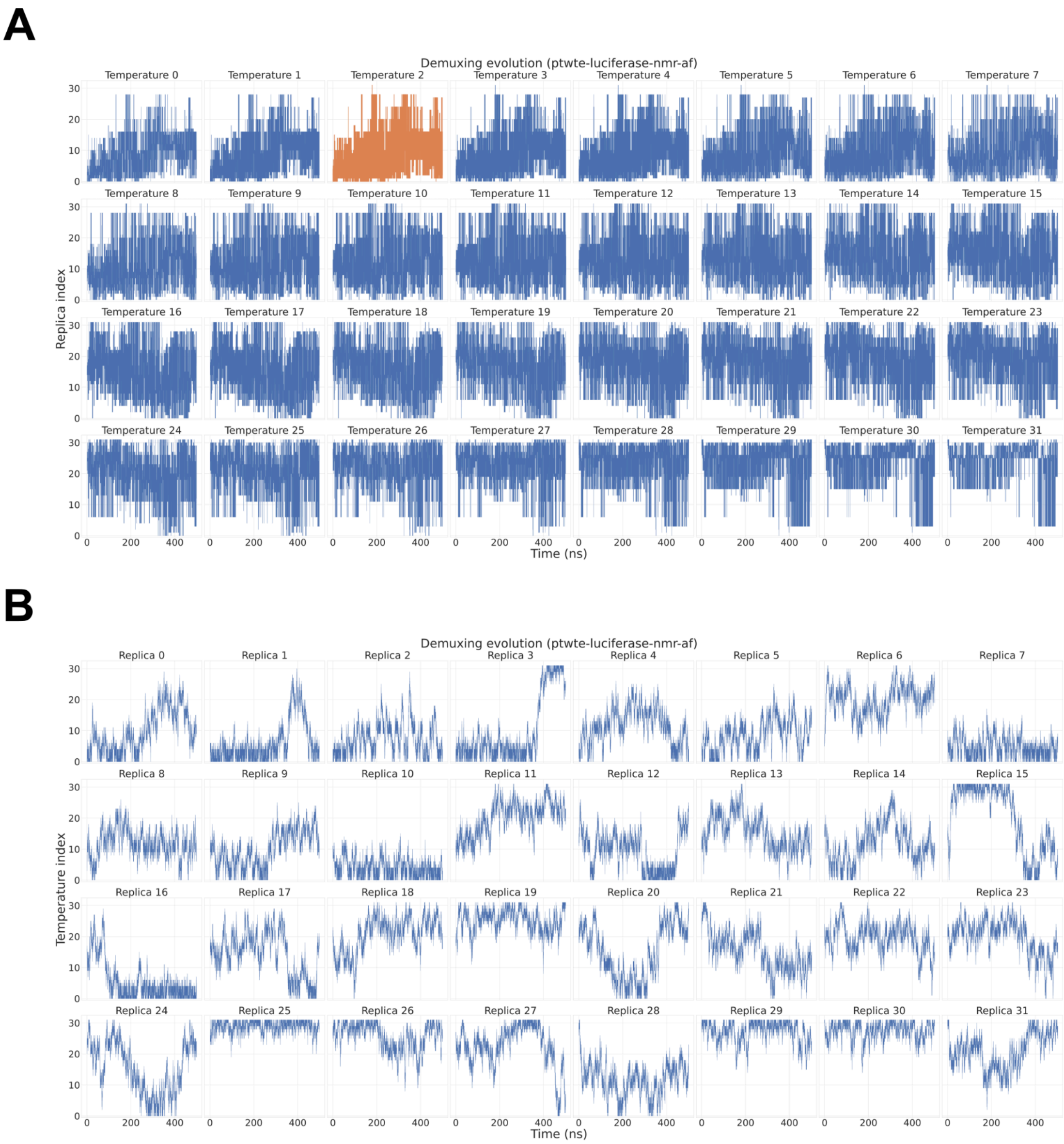


Figure S11. Demultiplexing analysis for the PT-WTE AF-NMR simulation. A) Analysis of the walk of each starting replica at each temperature index. As time progresses, we observe each replica being simulated at different temperatures. B) Analysis of the replicas that occupy each simulation temperature. Temperature 2, marked in orange, corresponds to the unbiased temperature used for analysis.

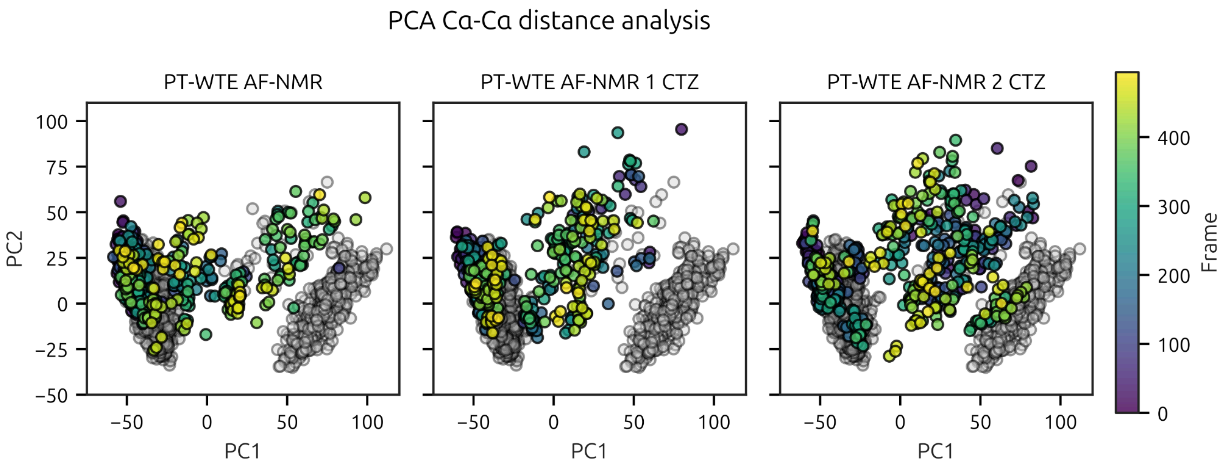


Figure S12: Time evolution of PT-WTE simulations projected on the PCA axis. Grey clusters represent AF (left) and NMR (right) simulation points. For further reference see Figure 2.

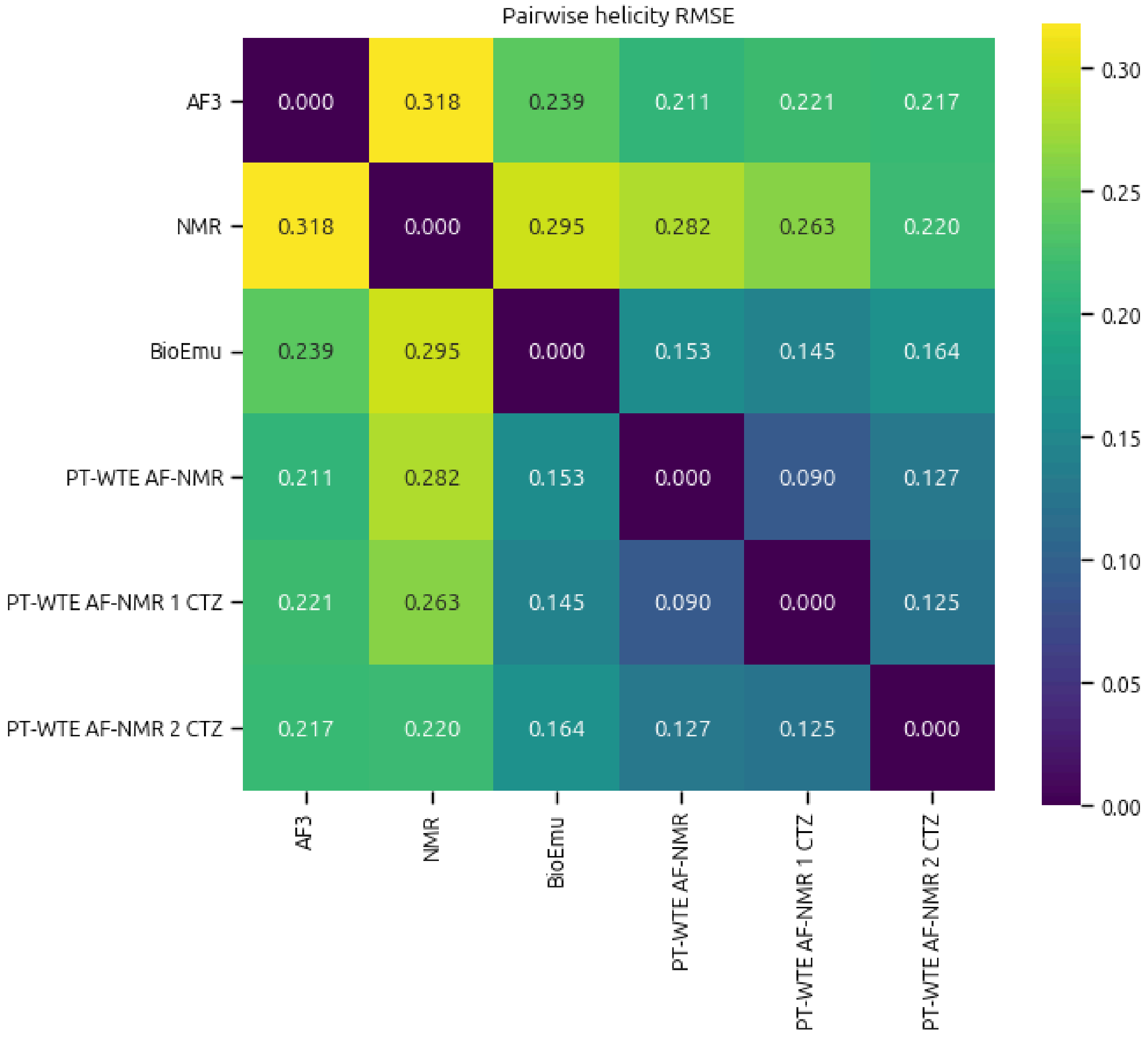


Figure S13: RMSE of the helicity profiles obtained for all simulations.

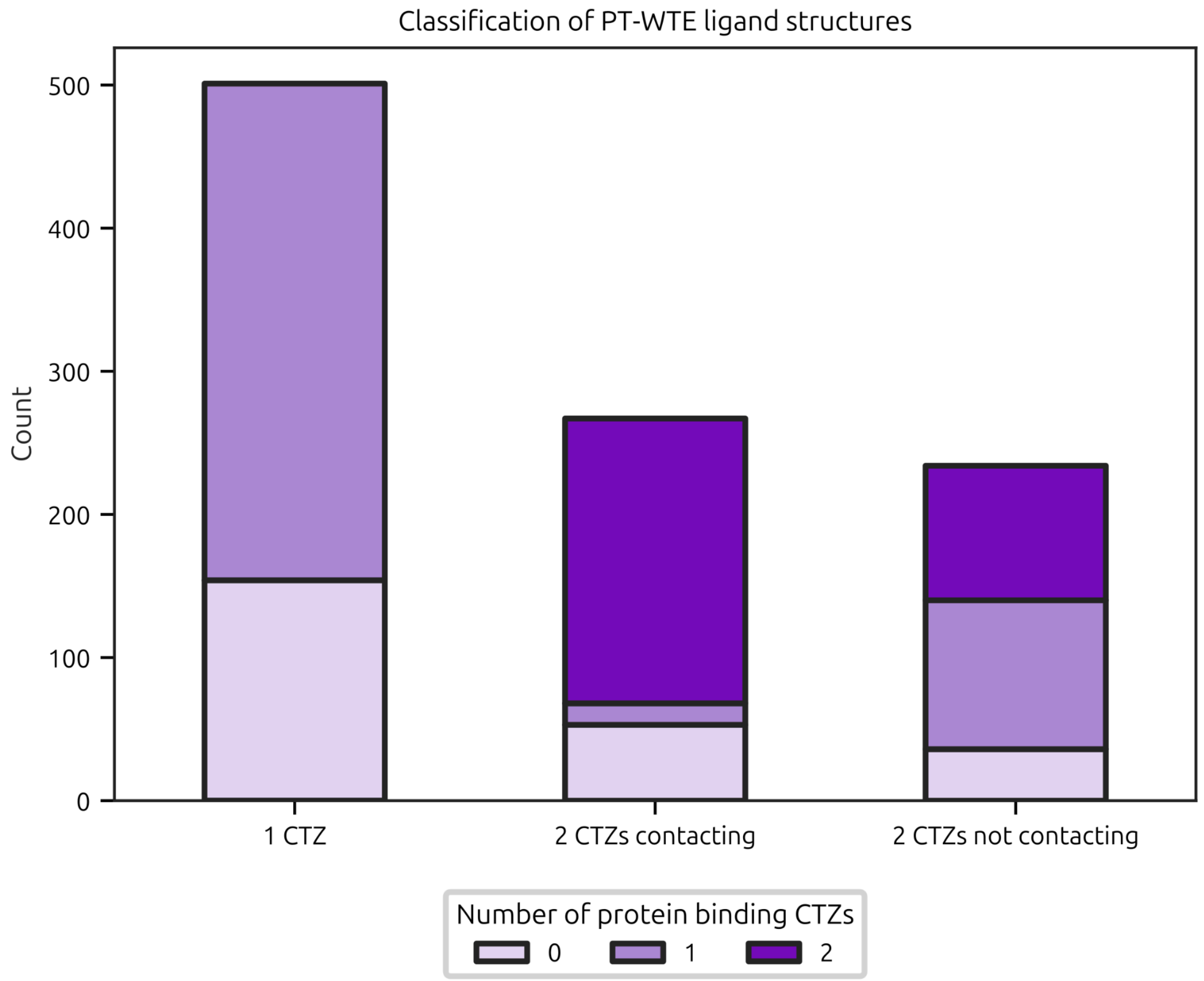


Figure S14: Classification of PT-WTE substrate simulations by the number of CTZs binding the protein and the presence of CTZ-CTZ aggregation.

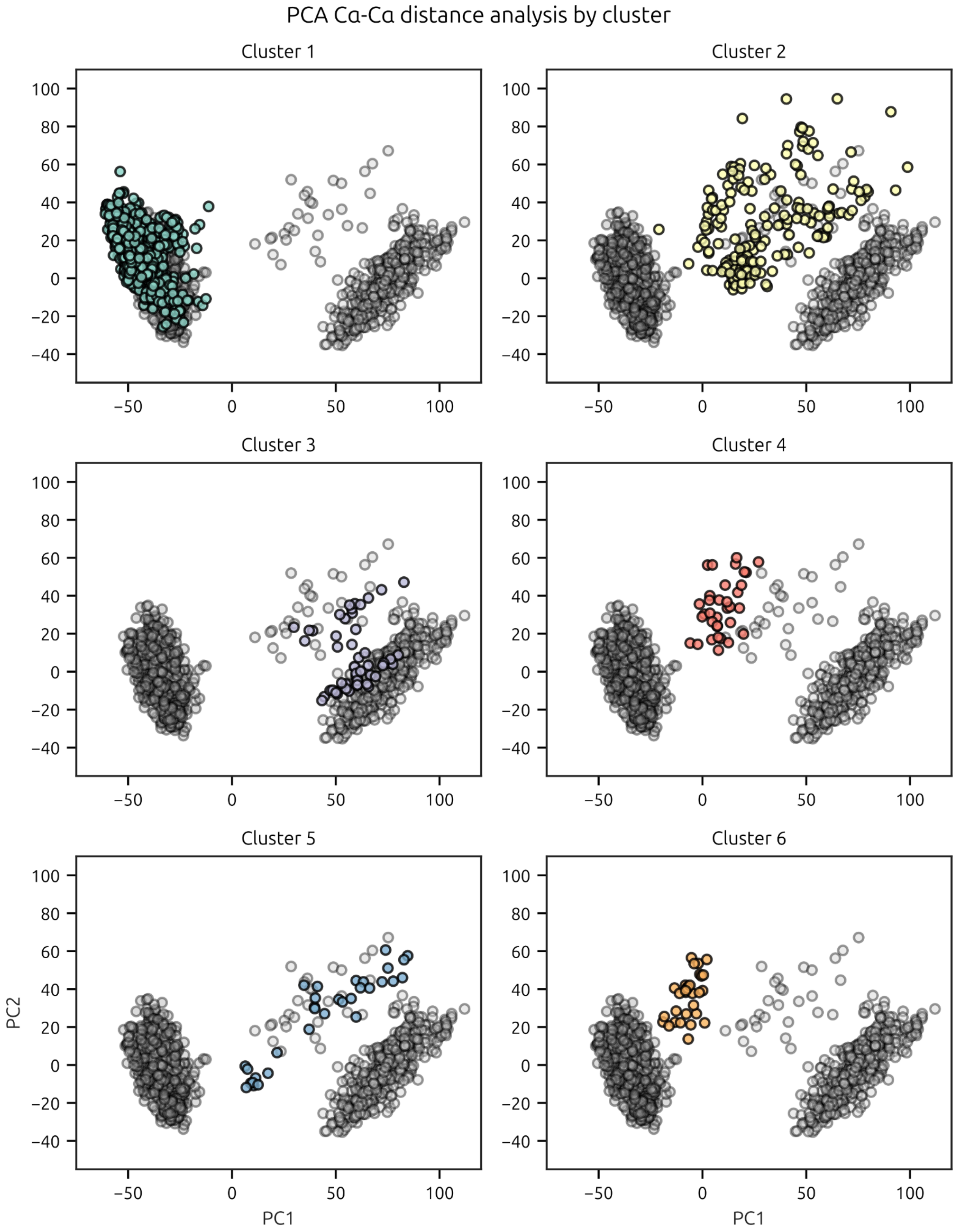


Figure S15: Projection of PT-WTE cluster structures in PCAs. The PCA space is the same as in Figure 2. Grey clusters represent AF (left) and NMR (right) simulation points.

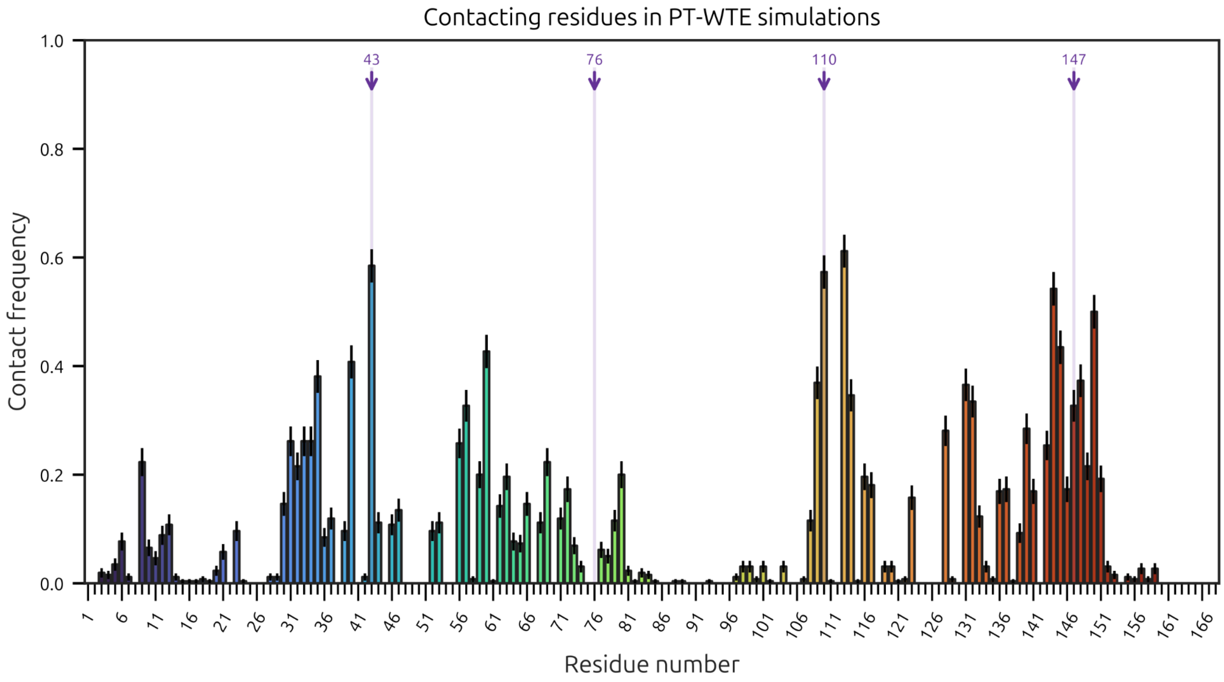


Figure S16: Frequency of contacting GLuc residues with CTZs from PT-WTE simulations with substrate. Results from cluster 1 and 2 were excluded to highlight contacts when CTZ is bound in a cavity and not the protein surface.

| Cluster | Number of CTZ in simulation | Population | Protein binding frames (%) | | CTZ-CTZ contacts (%) |
|---|---|---|---|---|---|
| | | | 2 CTZs | 1 CTZ | |
| **1** | 2 | 200 | 50 | 22 | 38 |
| | 1 | 298 | 58 | | - |
| | 0 | 339 | - | | - |
| **2** | 2 | 60 | 55 | 20 | 90 |
| | 1 | 45 | 84 | | - |
| | 0 | 78 | - | | - |
| **3** | 2 | 50 | 70 | 24 | 74 |
| **4** | 1 | 34 | 91 | | - |
| **5** | 2 | 32 | 38 | 56 | 0 |
| **6** | 2 | 31 | 84 | 13 | 94 |

Table S1: Contact and aggregation composition of every PT-WTE cluster.
